# Symmetric, explicit numerical integrator for molecular dynamics equations of motion with a generalized friction

**DOI:** 10.1101/172023

**Authors:** Ikuo Fukuda

## Abstract

A general mathematical scheme to construct symmetric, explicit numerical integrators of Newtonian equations of motion endowed with a generalized friction is provided for molecular dynamics (MD) study. The exact integrations are done for all the decomposed vector fields, including the one that contains the friction term. On the basis of the symmetric composition scheme with the adjoint for the resulting maps, integrators with any local order of accuracy can be systematically constructed. Among them, the second order P2S1 integrator gives the least evaluation of atomic force and potential, which are most time consuming in MD simulations. As examples of the friction function, three functional types are considered: constant, Laurent polynomial, and exponential with respect to the kinetic energy. Several MD equations of motion fall into these categories, and the numerical examinations of their integrators using a basic model system give positive results on the accuracy and efficiency. The extended phase-space scheme also presents an invariant function, which allows to easily detect numerical errors in the integration process by monitoring the function value.

## I. INTRODUCTION

Molecular dynamics (MD) [1–3] is a direct approach to investigate the characteristics of physical systems composed of atoms or molecules by assuming their interactions and numerically integrating a certainly defined equation of motion (EOM). A fundamental EOM is the Newtonian equation described by atomic coordinates *x* and velocities *v*, and it produces the micro-canonical ensemble with the constant energy. However, for conducting a realistic simulation that can mimic an experimental environment, it is desirable to produce a constant temperature and/or pressure.

For producing a constant temperature, Berendsen *et al.* [4] proposed a method combining the Newtonian EOM and an atomic-velocity scaling scheme. The Berendsen EOM is obtained by adding a force of the form –*g*_Bere_(*K*(*v*))*v* to the atomic force – ∇*V*(*x*) in the Newtonian EOM, where *V*(*x*) and *K*(*v*) is the atomic potential and kinetic energies, respectively. Namely, a friction is added and it is not a constant but dynamically changed according to the velocities *v*. This leads to the temperature control, where the instantaneous temperature and the kinetic energy *K*(*v*) is considered to be proportional. Despite the ambiguity of the resulting phase-space distribution [5], the Berendsen method is simple and stable, and it has been employed by many researchers [6] for e.g. biological simulations.

For producing a Boltzmann-Gibbs (BG) [or canonical] phase-space distribution 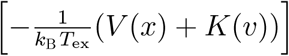 at the target temperature *T*_ex_, Nosé and Hoover proposed an alternative method [7, 8]. Their EOM, the Nosé-Hoover (NH) EOM, also employs dynamical friction term, but the frictional coefficient is no longer described by velocities *v* but by an additional dynamical variable *ζ*, such as –*g*_NH_(*ζ*)*v*. Here *ζ* is governed by its time derivative defined by the difference (up to a proportional constant) between the kinetic energy *K*(*v*) and the target temperature *T*_ex_: *ζ̇* = 2*K*(*v*) – *nk*_B_*T*_ex_. The NH has been widely used in MD simulations [9], and it also has provided a theoretical basis for developing general thermostatting method [10], sampling scheme [11], *ab initio* MD [12], and equilibrium and non equilibrium simulation techniques [13, 14].

For fluctuating the temperature of the BG distribution, the cNH has been proposed [15, 16]. The manner of the fluctuation of the temperature, 1/*β*, is not ad hoc, but completely governed by a (inverse-temperature) distribution function *f*(*β*). The EOM has a predetermined invariant density, which is a density of an invariant measure with respect to the Lebesgue measure, so that the probability density of realizing (*x*, *p*, *β*) is explicitly described. For this property, efficient sampling of space of (*x*, *p*) can be done by varying the temperature and by proceeded reweighting to a desired distribution, such as the BG distribution at the room temperature. By the temperature fluctuation, it may also be useful to observe the dynamics of the physical system in a non-equilibrium environment. The cNH EOM includes the frictional term taking the form of –*g*_cNH_ (*V*(*x*) + *K*(*v*),*ζ*)*v*.

The aim for these EOMs are not the same as stated, but their mathematical structure is the same such that they are described by an ordinary differential equation (ODE) of the form

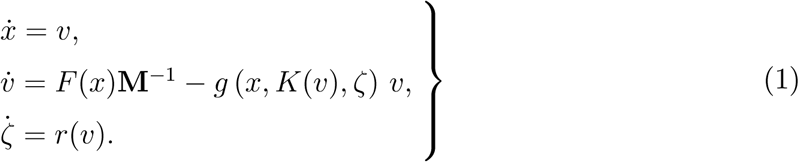

In this paper, a numerical integrator for this ODE, or a slightly generalized one, is developed. Since ODE (1) has a generalized friction that is a function of *K*(*v*) and extra variables, it is no longer (explicitly, at least) a Hamiltonian system [17, 18], so that the established scheme such as the symplectic integrators [19–24], including the Verlet method, cannot be straightforwardly used. Thus, the current integrator is constructed on the basis of the techniques previously developed for non-Hamiltonian systems [25]. Since the target of the current method is in the general form (1), it presents a unified integration scheme for the three kind of EOMs described above. For the NH EOM, since *g* has a simple structure such that *g*(*x*, *K*(*v*), *ζ*) = *g*_NH_ (*ζ*), its efficient integrators have been developed [26–29]. But this is a special case. In fact, integrators on the Berendsen EOM considered so far do not seem based on an established mathematical context, but based on heuristic approaches obtained by a combination of the leapfrog method and the velocity scaling. Regarding the cNH, its integrator has been constructed for the simplest case [15, 30]. Since the friction is related to the velocity *v*, it intimately concerns with the system temperature ∝ *K*(*v*) and the three EOMs relating to the temperature are the interesting examples. However, the target of ODE to construct integrators in this paper is in equation (1), which is not restricted for the applications to these three EOMs.

Note that regarding the EOM itself, not its numerical integrator, for a Newtonian EOM with a friction term, a number of mathematical considerations have been done. For example, stability [31, 32], long-time behavior [33–35], finite-time convergence [32, 36], control-theory algorithm [31], are investigated, and a friction in a general context, including subdifferentiated function [32, 34, 36] and a set-valued function [31], concerns to take into account e.g., a dry friction and a uncertain friction.

This paper provides an explicit, symmetric integrator for ODE (1). Symmetric property, or time-reversibility, which is one of the fundamental properties of ODEs, can be kept in the integration algorithm. In addition, by considering extended variable formalism, an invariant function for the extended ODE is provided. With the extended ODE, every solution in the original ODE, equation (1), remains unchanged. Thus, numerical error of the integration can be easily checked by monitoring the invariant function for the extended ODE, analogous to the Hamiltonian function for Hamiltonian ODE. By using symmetric composition technique [25, 37], the current method provides a systematic method so that higher order integrators can be systematically provided. To keep the less computational cost is also an important task for conducting MD simulations.

Section II specifically defines the target ODE in a general mathematical context and describes the NH, Berendsen, and the cNH EOMs as examples. Section III reviews the extended phase-space scheme, and Section IV gives the numerical integrator, where the vector field is decomposed into several components and each component is integrated exactly to make a fundamental integration map. The main result is on the integration of the component of the friction term, as detailed in Section IV A. A higher order method is stated in Section IV B, and an approximation method is provided in Section IV C for a practical convenience. As applications of the general scheme provided, in Section V, several kinds of friction functions are taken into account, and explicit integrator maps for the NH, Berendsen, modified Berendsen, and the cNH EOMs are presented. Section VI discusses the results of numerical simulations of these EOMs. Section VII gives the conclusion including remarks for future work.

## II. EQUATIONS OF MOTION

Our target EOM can be represented as the following ODE:

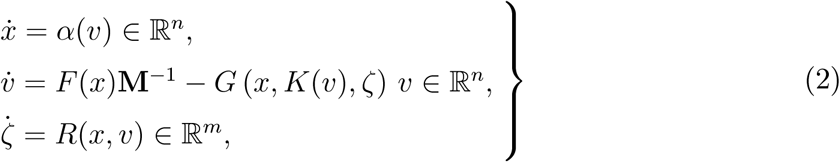

where *x* ≡ (*x*_1_, …, *x_n_*) ∈ *D* ⊂ ℝ^*n*^, *v* ≡ (*v*_1_, …, *v_n_*) ∈ *B* ⊂ ℝ^*n*^, *F*(*x*) ∈ ℝ^*n*^, and *K*(*v*) ∈ ℝ can be interpreted as the atomic coordinates, velocities, force (smooth vector-valued function on a domain *D*), and kinetic energy, respectively, of a physical system of *n* degrees of freedoms. Here *ζ* ∈ ℝ^*m*^ is a kind of extended variable, as detailed below. Thus a phase space point is *ω* = (*x*, *p*, *ζ*), which becomes an element of the target phase space Ω: = *D* × *B* × ℝ^*m*^ ⊂ ℝ^*N*^ with *N* ≡ 2*n* + *m*. Mathematical conditions for the three functions, *α*, *G*, and *R*, are as follows:

### Condition 1

*F*: *D* → ℝ^*n*^, *α*: *B* → ℝ^*n*^, *G*: *D* × *B_K_* × ℝ^*m*^ → ℝ, *and R*: *D* × *B* → ℝ^*m*^, *are of class C^r^* (*r* ≥ 1), *and K*: *B* → ℝ, *v* ↦ (*v*|*vM*)/2. *Here*, *B and B_K_ are (non*-*empty) open sets of* ℝ^*n*^ *and* ℝ, *respectively*, *satisfying K*(*B*) ⊂ *B_K_ and a normal property*: λ*v* ∈ *B for any v* ∈ *B and* λ > 0. *The mass parameters* M *is a symmetric*, *positive definite square matrix of size n (over* ℝ).

We give examples that falls into the form of equation (2):

### Example 2

*α*(*v*) ≡ *v* ∈ *B* ≡ ℝ^*n*^, *G* (*x*,*k*,*ζ*) ≡ *ζ*/*Q with B_K_* ≡ ℝ, *and R*(*x*,*v*) ≡ 2*K*(*v*) – *nk_B_T*_ex_ *with m* ≡ 1, *where Q* > 0 *is a temperature*-*control constant*, *T*_ex_ > 0 *a target temperature*, *and k_B_ Boltzmann’s constant*, *give the NH equation [7*, *8]:*

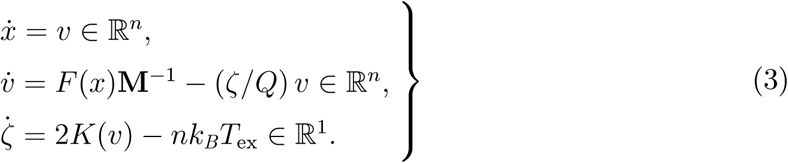

### Example 3

*α*(*v*) ≡ *v* ∈ 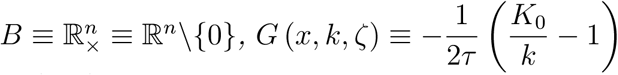 *with B_K_* ≡ ℝ_+_ ≡ (0,∞), *and formally set R*(*x*,*v*) ≡ 0, *where K*_0_ > 0 *is a target kinetic energy value and τ* > 0 *is the temperature*-*control time constant*, *give the Berendsen EOM [4]:*

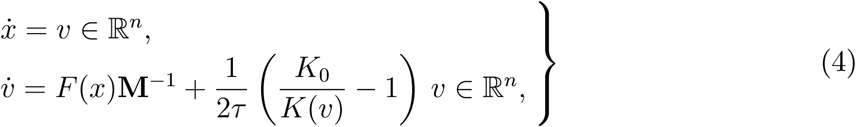

*where we have omitted the decoupled EOM*, *ζ͘* = 0. *Condition 1 can be met.*

### Example 4

*Coupled NH (cNH) equations of motion is introduced to fluctuate the temperature of the heat bath for the physical system [15*, *30]. The cNH couples two NH equations*, *where one is the NH equation of the physical system represented by* (*x*,*p*, *ζ*) *and the other one is the NH equation of the temperature system represented by* (*Q*, 𝓟, *η*). *The cNH is represented by*

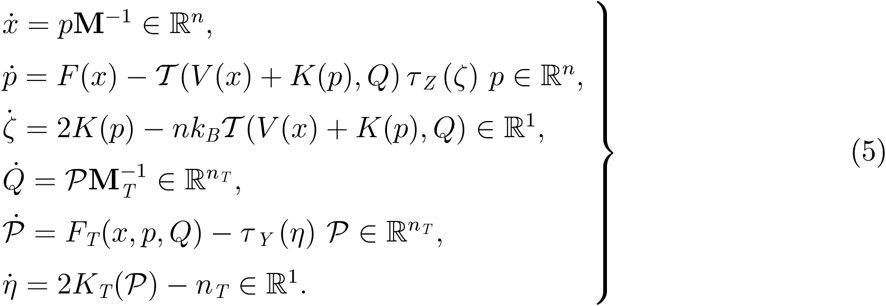

*The vector field can be split into two vector fields*, *and the problem is reduced to seek an explicit solvable decompositions of the two fields*, *one of which yields an ODE*,

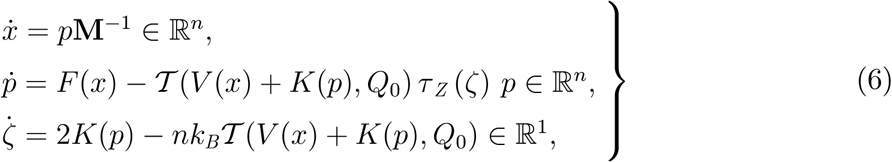

*and the other yields an ODE*,

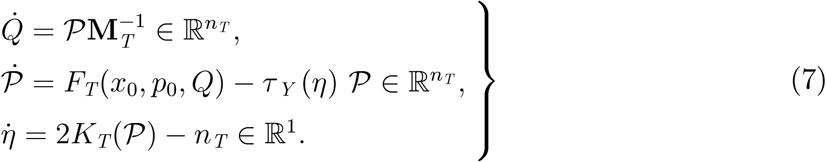

*Equation (6) can be represented as a form of (2) by substitutions of v* ≡ *p* ∈ *B* ≡ ℝ^*n*^, *α*(*p*) ≡ *p***M**^−1^, *G* (*x*,*k*,*ζ*) = 𝓣(*V*(*x*) + *k*,*Q*_0_) *τ_Z_* (*ζ*), *and R*(*x*,*p*) ≡ 2*K*(*p*) – *nk_B_*𝓣(*V*(*x*) + *K*(*p*),*Q*_0_), *where Q*_0_ *corresponding to an initial value of Q. τ_Z_*: ℝ→ ℝ *and* 𝓣: ℝ × ℝ*^n_T_^* → ℝ *are C^r^ functions and B_K_* ≡ ℝ. *Similarly*, *equation (7) is reduced to (2) by α*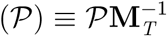 *with* 𝓟 ∈ *B* ≡ ℝ^*n*_T_^ *(n_T_* ∈ ℕ), *G* (*Q*, *k*, *η*) ≡ *τ_Y_* (*η*) *with B_K_* ≡ ℝ, *and R*(*Q*, 𝓟) = 2*K_T_*(𝓟) – *n_T_* ≡ 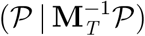 – *n_T_*, *with using x*_0_,*p*_0_ *that correspond to initial values of x*,*p*. *F_T_ and τ_Y_ are C^r^ functions. Formal redefinitions of masses*, **M** *and* **M**_*T*_, *and forces*, *F and F_T_*, *to conform to the form of equation (2) are clear. See Example 8 for more details including the specific forms of the functions used here.*

### Remark 5

*We have a slight extension in which α*(*v*, *ζ*) *and F*(*x*, *ζ*) *can be used*, *instead of α*(*v*) *and F*(*x*), *respectively. Namely*, *α and F allow dependencies of ζ*, *wherein α*: *B* × ℝ^*m*^ → ℝ^*n*^ *and F*: *D* × ℝ^*m*^ → ℝ^*n*^ *should be of class C^r^. The method stated below can also be applied for this extension.*

## III. EXTENDED ODE WITH INVARIANT FUNCTION

The target ODE (2) can be generally represented as

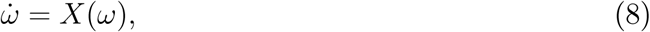

where *X* becomes a smooth vector field defined on an open set Ω of ℝ^*N*^. This ODE does not necessarily have an invariant function, such as the Hamiltonian function of a Hamiltonian system, so that the confirmation of accuracy in the numerical integration may not be done by simply monitoring the invariant-function value but require other efforts.

This inconvenience can be overcome by considering an extended system [25] for the target system. The extended system is defined by associating an additional variable v ∈ ℝ to the original variables *ω* ≡ (*ω*_1_, …, *ω_N_*) ∈ Ω and represent them by *ω*′ = (*ω*, v) as a point of an “extended space” Ω′ ≡ Ω × ℝ. The extended ODE [25] on Ω′,

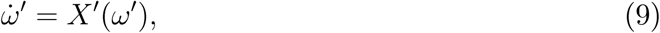

is defined by

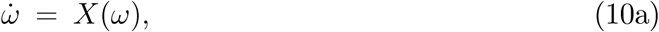

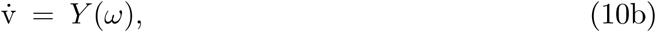

where

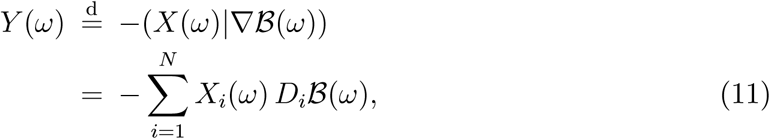

with 𝓑 being an arbitrary smooth function on Ω. Then it is straightforwardly seen that a function

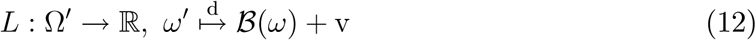

becomes an invariant of (9); i.e.,

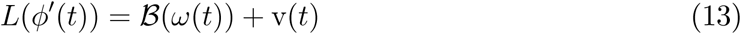

is constant for any time *t* for an arbitrary solution *ϕ*′ = (*ω*, v) of (9). Thus, by monitoring the value of *L*(*ϕ*′(*t*)) while numerically integrating (9), we can check the numerical error. It should be noted that every solution, *t* ↦ *ω*(*t*), in the original ODE (10a) is not perturbed at all by attaching (10b). The multiple extended-variable formalism, where v ∈ ℝ^*l*^ is generally considered, is useful to enhance the handling in actual simulation [29].

There are many choices of the function 𝓑. Several examples of 𝓑 and the resulting *Y* are presented below taking the EOMs introduced in section II. Assume the existence of the potential function *V* such that *F* = – ∇*V*.

### Example 6

*For the NH equation*,

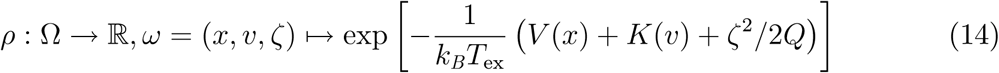

*becomes an invariant density*, *viz.*, div *ρX* = 0. *In such a case*, *a standard manner [25] can be chosen in such a way that* 𝓑 = –*c* ln *ρ with an arbitrary constant c*, *which yields*

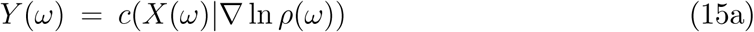

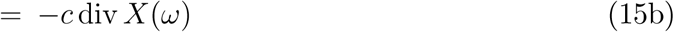

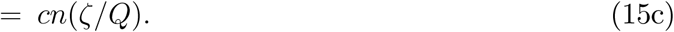

*By putting c* ≡ *k_B_T*_ex_, *we have*

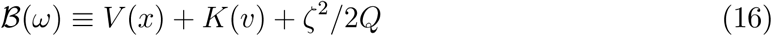

*and the invariant*

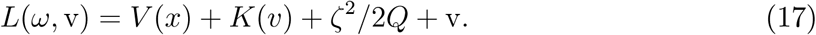

### Example 7

*Although the existence of an invariant density is unknown for the Berendsen EOM*, *remind that* 𝓑 *can be chosen freely. One of the natural choices of* 𝓑 *that are analogous to equation (16) is the total energy*, *viz*.,

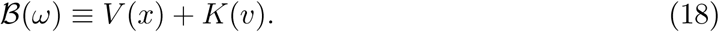

*We then have*

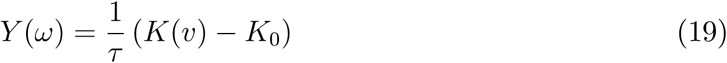

*and*

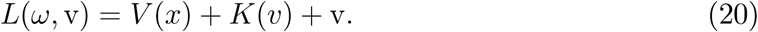

*It is a trivial note that by considering R*(*x*, *v*) ≡ 0 *expressed in Example 3*, 𝓑 *can be taken as* 𝓑(*x*,*v*) ≡ *V*(*x*) + *K*(*v*) + *G*(*ζ*) *for any smooth G*, *so that L has the same form as the NH: L*(*ω*,v) = *V*(*x*) + *K*(*p*) + *G*(*ζ*) + v, *where ζ is just a constant variable.*

### Example 8

*The cNH has an invariant density ρ [15*, *30] defined by*

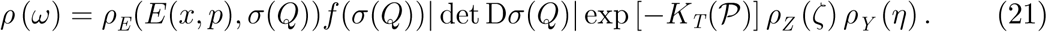

*By realization of this density in the dynamics*, *the original physical system represented by* (*x*,*p*) *obeys the distribution density ρ_E_*(*E*(*x*,*p*), *σ*(*Q*)) *having an inverse temperature β* ≡ *σ*(*Q*) ∈ ℝ_+_, *which also fluctuates according to the distribution density f* (*β*). *For instance*, *the choice*

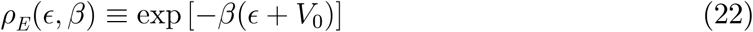

*with n_T_* ≡ 1 *corresponds to the situation in which the inverse temperature β of the heat bath*, *which is characterized by the BG distribution as usual*, *fluctuates (in other words*, *an automatic continuous temperature change in the replica exchange method can be realized). According to the change of the dynamical variable Q in the dynamics*, *β* ≡ *σ*(*Q*) *varies through a suitable smooth function σ. σ and f can be freely chosen under certain mathematical conditions described in [15*, *30]. A typical choice of ρ_Z_ and ρ_Y_ are the Gaussian functions such as*

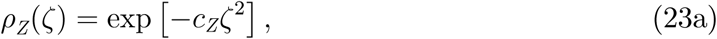

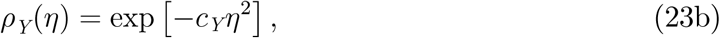

*and then *τ*_Z_* (*ζ*) = 2*k_B_c_Z_ζ and τ_Y_* (*η*) = 2*c_Y_η in equation (5). Now*, *regarding the choice of* 𝓑 *for the cNH*, *the same procedure as the NH can be used*, *due to the existence of the invariant density (21)*. *If we set* 𝓑 ≡ – ln *ρ*, *then*

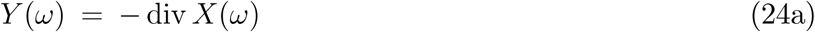

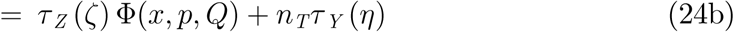

*with*

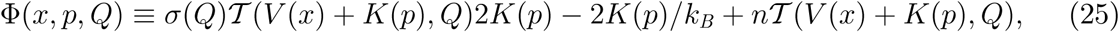

*and the invariant is L*(*ω*, v) = – ln *ρ*(*ω*) + v, *which becomes*

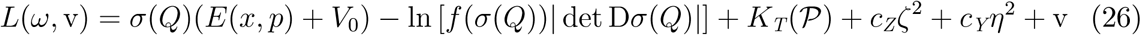

*if we use equations (22) and (23). Note that* 𝓣(*V*(*x*) + *K*(*p*),*Q*) *in both equation (25) and original EOM (5) is derived from ρ_E_ [30]. The explicit form of* 𝓣 *in case of density (22) is given as*

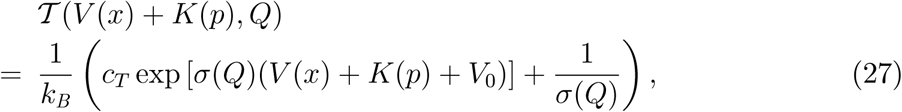

*where c_T_* ≥ 0 *is a constant [30].*

## IV. INTEGRATOR

### A. First-order Integrator

For constructing an integrator, we first decompose a target vector field *X*′ and then compose the corresponding extended-phase-space maps [25]. In the current case, we give
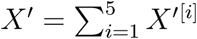

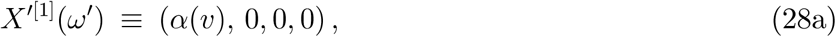

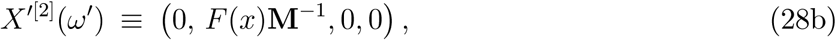

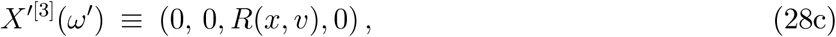

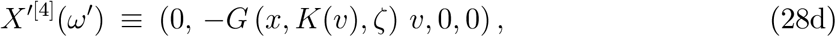

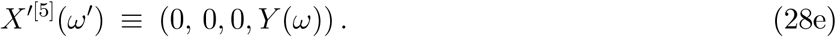

For an individual vector field *X*′^[*i*]^, the flow Φ^[*i*]^, for which *t* ↦ Φ^[*i*]^(*ω′*; *t*) denotes the solution of *ω̇*′ = *X*′^[*i*]^ (*ω′*) with an initial value *ω′* = (*x*;*p*; *ζ*; *v*) ∈ Ω′, can be defined as an extended-phase-space map: Φ^[*i*]^: Ω′ × ℝ → ℝ^*N′*^, where *N′* ≡ *N* + 1. Except *i* = 4, we easily see that Φ^[*i*]^ is explicitly represented by

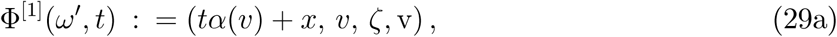

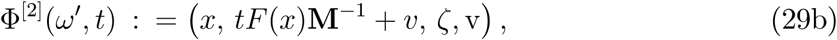

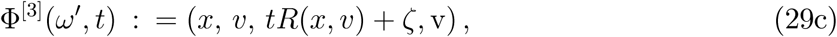

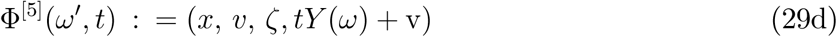

for all *t* ∈ ℝ. To be precise, an exact flow takes values in Ω′, so that the domain of definition of Φ^[*i*]^ should be generally restricted into Ω′^[*i*]^ = (Φ^[*i*]^)^−1^(Ω′) since the extended phase space Ω′ is not necessarily the whole ℝ^*N′*^ but is *D* × *B* × ℝ^*m*^ × ℝ (exceptionally, Ω′^[*i*]^ = Ω′ × ℝ for *i* = 3, 5). Note that the decomposition of *X′* is natural in that the additional quantities v and *Y* do not affect the solutions of a decomposed original ODE *ω̇* = *X*^[*i*]^(*ω*) for *i* = 1, 2, and 3, where *X*^[*i*]^ is obtained by the same type of the decomposition of the original field
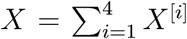
(viz., *X*^[*i*]^ is defined by removing 0 in the last column of (28a), (28b), and (28c) for *i* = 1, 2, and 3, respectively).

To get the explicit form of Φ^[4]^ seems nontrivial, but it can be done. It is the main task of this work to represent explicitly Φ^[4]^ and construct efficient integrator for ODE (2). Before obtaining the solution of

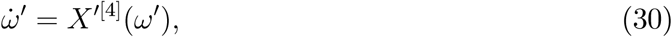

we prepare two lemmas. The first one is an issue of an elementary 1-dimensional ODE but needed to consider the domain of definition of integrator maps with care.

#### Lemma 9

*Let* λ: ℝ ⊃ *U* → ℝ_×_ ≡ ℝ ∖ {0} *be of class C^s^ (s* ≥ 0), *with U being an open interval. Then the solution φ of an ODE*,

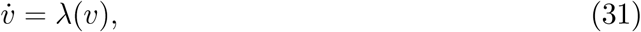

*with an initial condition* (*t*_0_, *v*_0_) ∈ ℝ × *U is given by*

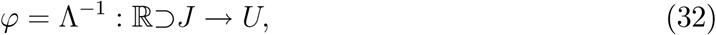

*where*

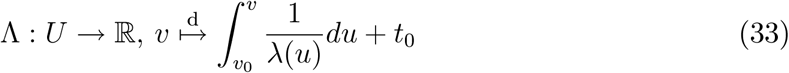

*being a strictly monotonic*, *C*^*s*+1^ -*diffeomorphic function*, *and J* ≡ Λ(*U*) *being an open interval including t*_0_.

#### Proof.

The assumptions for λ and *U* ensure that λ has a definite signature on the whole *U*, via the intermediate value theorem. Thus Λ is well defined and strictly monotonic so that it is 1-1, as well as of class *C*^*s*+1^. We see that *D*Λ(*v*): ℝ → ℝ, *y* ↦ (1/λ(*v*))*y*, is linear isomorphic and homeomorphic for any *v* ∈ *U*. Thus, *J* = Λ(*U*) is an open set in ℝ, and Λ: *U* → *J* = Λ(*U*) becomes *C*^*s*+1^ diffeomorphism (via the inverse function theorem). Since *U* is connected and Λ is continuous, *J* becomes an interval. We also see that ∀*t* ∈ *J*, ∃*D*Λ^−1^ (*t*) = (*D*Λ(Λ^−1^(*t*)))^-1^ = λ(Λ^−1^(*t*)), implying *φ* = Λ^−1^ is a solution of ODE (31). Noting *J* ∋ Λ(*v*_0_) = *t*_0_, we have the initial condition *φ*(*t*_0_) = Λ^−1^(*t*_0_) = *v*_0_. ◼

The uniqueness of the solution for the initial value problem of equation (31) is clear if *s* ≥ 1, and actually the uniqueness holds even for *s* = 0.

Due to Condition 1, we have the following Lemma regarding *B_K_*:

#### Lemma 10

∀(*x*,*ζ*) ∈ *D* × ℝ^*m*^, *there exist at most countable disjoint open subsets U_α_* ⊂ ℝ *(α* ∈ *A) such that*

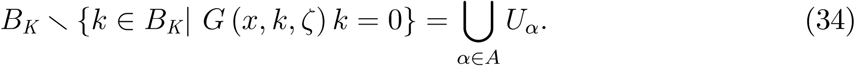

*Here*, *every U_α_ (α* ∈ *A) becomes an interval if* ∃*k* ∈ *B_k_*, *G*(*x*,*k*,*ζ*) ≠ 0 *and U_α_* = ∅ *otherwise. For any α* ∈ *A*, 0 < ∃*k* ∈ *U_α_ implies U_α_* ⊂ (0,∞).

#### Proof.

Fix an arbitrary (*x*,*ζ*) ∈ *D* × ℝ^*m*^. Assume the case in which ∃*k* ∈ *B_k_*, *G*(*x*,*k*,*ζ*) ≠ 0. The continuity of *G* indicates that *μ*: *B_K_* → ℝ, 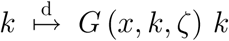 is *C*^0^ and thus *U* ≡ *B_K_* ∖ Ker *μ* is an open set in *B_K_* and is open also in ℝ since *B_K_* is so. If we can say *U* is not empty, then we have the expression of (34). Let *U* be empty. Then ∀*k* ∈ *B_K_* ∖ {0}, *G* (*x*, *k*, *ζ*) = 0. Here *B_K_* ∖ {0} is not empty since the existence of nonzero *b* ∈ *B* implies 0 ≠ *K*(*b*) ∈ *K*(*B*) ⊂ *B_k_*. Thus, it follows from the continuity of *G* that *G* (*x*, 0, *ζ*) = 0 if 0 ∈ *B_K_*. We have so *G* (*x*,*k*,*ζ*) = 0 for all *k* ∈ *B_K_*, which contradicts the assumption of *G* in the current case. To show the remaining issue, suppose ∃*k*_+_, *k*_−_ ∈ *U_α_* such that *k*_+_ > 0 and *k*_−_ ≤ 0. This implies 0 ∈ [*k*_−_, *k*_+_] ⊂ *U_α_,* which contradicts 0 ∈ Ker*μ*. Next, in case of ∀ ∈ *B_K_ G* (*x*, *k*, *ζ*) = 0, the LHS of (34) is empty, so that *U_α_* can be set as ∅ and the last issue also trivially holds. ◼

Now we have the explicit form of the solution of the target ODE as follows:

#### Proposition 11

*A solution of equation (30) with initial value* (*x*_0_, *v*_0_, *ζ*_0_, *v*_0_) ∈ Ω′ *is uniquely given by* ℝ ⊃ *J* → ℝ^*N′*^, *t* ↦ (*x*_0_,*v*(*t*),*ζ*_0_, *v*_0_), *where v is defined as follows: (i) if* ∃*α* ∈ *A such that k*_0_ ≡ *K*(*v*_0_) ∈ *U_α_*, *then*

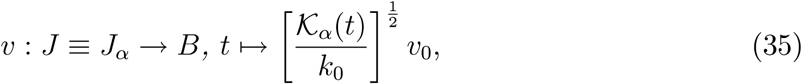

*which is defined by*

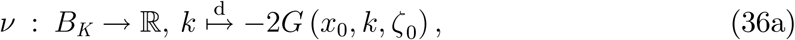

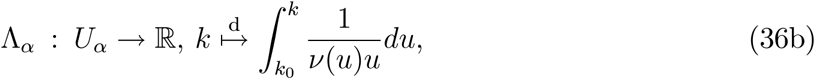

*with J_α_* ≡ Λ_*α*_ (*U_α_*) *being an open interval including* 0, *and*

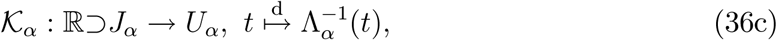

*which becomes C*^*r*+1^ -*diffeomorphic; (ii) if G* (*x*_0_, *k*_0_, *ζ*_0_) *k*_0_ = 0, *then v*: *J* ≡ ℝ → *B*, *t* ↦ *v*_0_, *which is a constant map. Here*, *in general*, *A and n depend on* (*x*_0_, *ζ*_0_), *and U_α_*, Λ_α_, *J*_α_, *and* 𝓚_*α*_ *depend on* (*x*_0_,*k*_0_,*ζ*_0_).

#### Proof.

Fix any (*x*_0_, *ζ*_0_) ∈ *D* × ℝ^*m*^ and *v*_0_ ∈ ℝ, and note *k*_0_ = *K*(*v*_0_) ∈ *K*(*B*) ⊂ *B_K_*. According to *G* and (*x*_0_, *ζ*_0_), we encounter two cases: ∀*k* ∈ *B_K_ G* (*x*_0_,*k*,*ζ*_0_) = 0 or not. For the former case, (ii) applies for any *k*_0_. For the latter case, Lemma 10 indicates *B_K_* = {*k* ∈ *B_K_*| *G* (*x*_0_,*k*, *ζ*_0_) *k* = 0} ∪ ⋃_*α*∈*A*_ *U_α_*, where every *U_α_,* depending on (*x*_0_,*ζ*_0_), is an open interval. Thus, according to *k*_0_, Case (i) applies if ∃*α* ∈ *A*,*k*_0_ ∈ *U_α_*, and Case (ii) applies otherwise. Since Case (ii) is obvious, we prove Case (i) blow.

Let ∃*α* ∈ *A* such that *k*_0_ ∈ *U_α_*, which also implies that the selected *U_α_* depends on *k*_0_ as well as (*x*_0_, *ζ*_0_). Define a *C^r^* line field λ: *U_α_* → ℝ_×_, 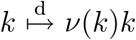 (*x*_0_, *k*, *ζ*_0_) *k*, and apply Lemma 9 to consider an ODE

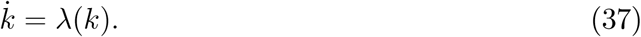

Then we see that its solution 𝓚_*α*_ with an initial condition (0, *k*_0_) ∈ ℝ × *U_α_* is given as equations (36a)–(36c), with Λ_*α*_ being *C*^*r*+1^-diffeomorphic and *J* = *J_α_* being an open interval including *t*_0_ = 0.

Now, we see that

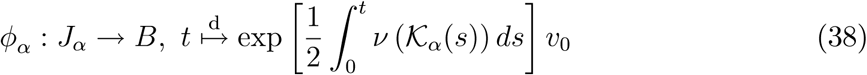

is well defined, where *ϕ_α_* (*t*) ∈ *B* comes from the normal property stated in Condition 1. It follows that *ϕ_α_*(0) = *v*_0_ and

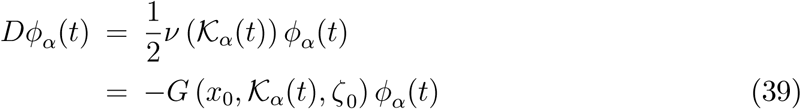

for any *t* ∈ *J_α_.* Thus, if

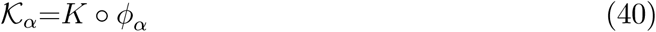

holds true, then the uniqueness of the solution of the initial value problem of ODE (30) (notice *X*′^[4]^ is of *C^r^* with *r* ≥ 1)indicates that *J_α_* → Ω′, *t* ↦ (*x*_0_, *ϕ_α_*(*t*), (*ζ*_0_, v_0_), becomes the solution sought for.

To show equation (40), first note that for any *t* ∈ *J_α_*, there hold equalities

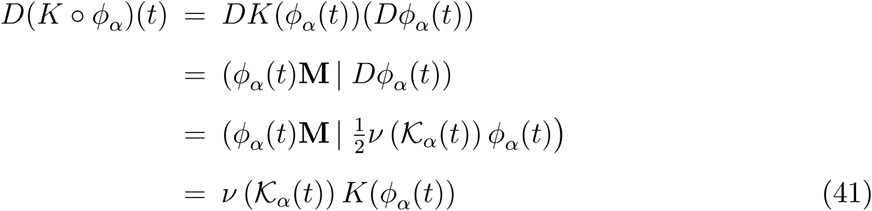

and hold an inequality *ϕ_α_*(*t*) ≠ 0 (since *v*_0_ ≠ 0), which yields *K*(*ϕ_α_*(*t*)) > 0. Thus

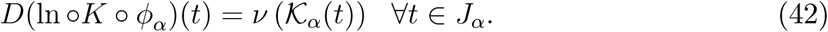

We see ln ∘𝓚_*α*_: *J_α_* → *U_α_* → ℝ is well defined since *U_α_* ⊂ (0,∞), which comes from Lemma 10 and 0 < *k*_0_ ∈ *U_α_.* Thus, the fact that 𝓚_*α*_: *J_α_* → *U_α_* is a solution of ODE (37) leads to

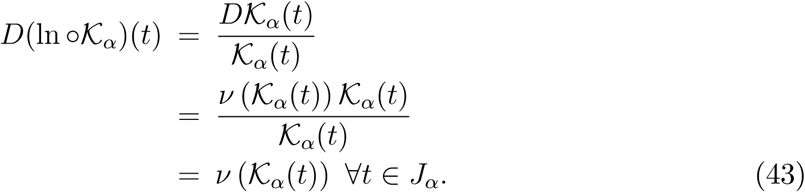

Equations (42) and (43) and the fact that *J_α_* is a domain imply

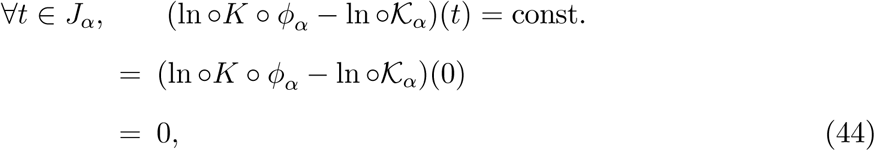

where the last line is due to *K*(*ϕ_α_*(0)) = *K*(*v*_0_) = *k*_0_ = 𝓚_*α*_(0). Hence we have proved equation (40).

Finally we show that *ϕ_α_* is equal to *v* defined by equation (35). Using equation (43), for any *t* ∈ *J_α_*, we get

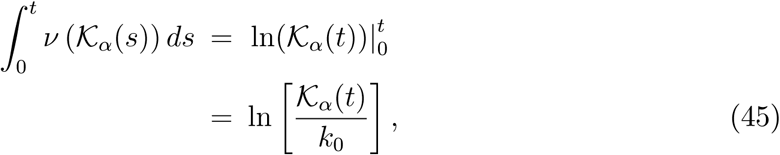

so that

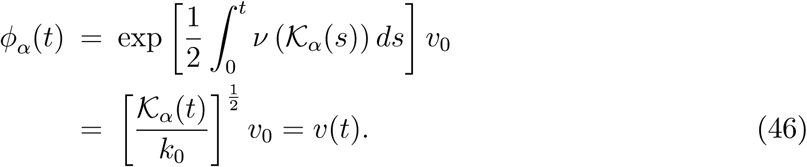

Namely, we have obtained *ϕ_α_* = *v* with *𝓚_α_*=*K* o *v* on *J_α_*. The dependence of Λ_α_ on (*x*_0_, ζ_0_) comes from that of *U_α_* and *v*, and the dependence on *k*_0_ comes from both *U_α_*and the lower bound of the integral. The dependence of *J_α_* and *𝓚_α_* on (*x*_0_, *k*_0_, ζ_0_) follows from that of *U_α_* and Λ_α_. We thus completed the proof. ◼

Hence we have the explicit form of Φ ^[4]^ as

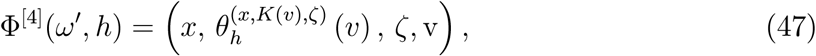

where

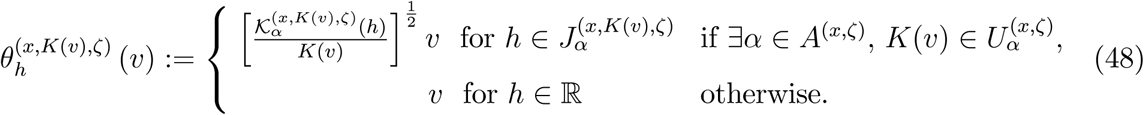

Here
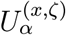
≡ *U_α_* and *A*^(*x*,*ζ*)^ ≡ *A* denote explicitly the dependence of *x* and *ζ*, and similarly
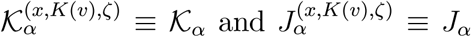
denote the dependence of *x*, *K*(*v*), and *ζ*. However, since equation (48) seems complicated, we consider a simpler expression for convenience. First, to make a unified expression, we introduce a special index value α = 0 and

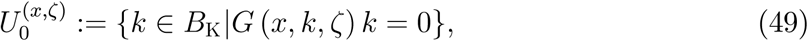

so that any *K*(*v*) belongs to
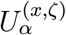
for a certain *α* ∈
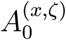
≡ *A*^(*x*,*ζ*)^∪{0}. Since this *α* is uniquely determined by *K*(*v*), we can denote *α* = [*K*(*v*)] (in fact, [⋅] can be viewed as the natural surjection from *B_K_* to a quotient space
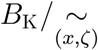
generated by an equivalent relation such that
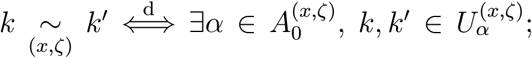
then we have a natural identification between
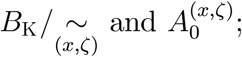
note that
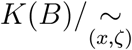 also arrises, which corresponds to a classification of the kinetic-energy space *K*(*B*), and also a velocity-space classification
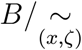
arrises). For this reason, to remove the redundancy in the notation
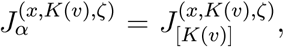
we can rewrite it as *J*^(*x*,*K*(*v*),*ζ*)^, or more simply, *J^ω^*, for *ω* = (*x*,*v*, *ζC*). Similarly,*𝓚^ω^* can be used instead of
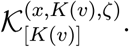 Second, define

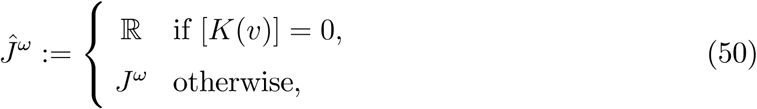

and

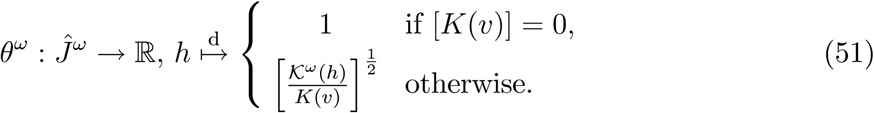

Then we finally have a compact form as

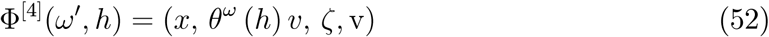

using simple expressions (50) and (51). It should be pointed out that, first, to obtain *θ^ω^* (*h*) by equation (51), a switching we should care is only [*K*(*v*)] = 0 or not, where [*K*(*v*)] = 0 is equivalent to *G* (*x*,*K*(*v*).*ζ*) *K*(*v*) = 0. Second, the dependence of *𝓚^ω^* on *ω*, accurately, on *x*, *K*(*v*) and *ζ*, is through both the domain of the definition and the range of 𝓚^*ω*^, in general. So that although the behavior of 𝓚^*ω*^ depends on these quantities, the function form of 𝓚^*ω*^does not depend on these quantities (the indefinite integral in equation (36b) does not depend on these). Third, the switching is not a discontinuous one (in fact, Φ^[4]^ is of *C^r^*), and the constant function *θ^ω^* = 1 in the case [*K*(*v*)] = 0 can be viewed as a limiting case of *G*(*x*,*K*(*v*),*ζ*) *K*(*v*) → 0 since the inverse function of a rapidly growing function Λ_*α*_[equation (36b)] due to a large
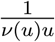
becomes a slowly growing function and tends to the constant. Last, Φ^[4]^ is not affected by *v* and *Y*, as well as the other Φ^[*i*]^ (*i* = 1. 2. 3), as noted in section IV A. Namely, (*ω*,*h*) ↦ (*x*, *θ^ω^* (*h*) *v*. *ζ*) is obtained as the exact flow of *X*^[4]^, which is defined by removing 0 in the last column of (28d).

Therefore, combining equation (52) with equation (29), we get a first-order integrator,

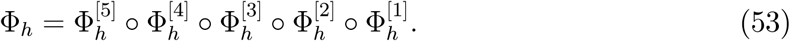

Here, in general, for maps Φ^[*i*]^: Ω′ × ℝ *O*′^[*i*]^ → ℝ^N^′ and time *h* ∈ ℝ, we can define local phase-space maps
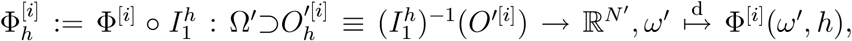
where
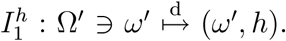
Equation (53) is thus obtained for Φ^[*i*]^ (*i* = 1.… . 5) defined by equation (52) with *O*′^[4]^ = ⋃_(*ω*>,*v*)∈*Ω*_′ {(*ω*, *v*)} × *ª̂^ω^* for *i* = 4 and equation (29) with *O*′^[*i*]^ = Ω′ × ℝ for *i* ≠ 4. Note that if every map has the form of
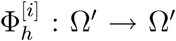
then their composition can be freely done, but
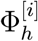
(*i* = 1. 2) (as noted in section IV A) and
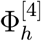 do not have this form in general. Thus we shall assume hereafter that the domain of definition of the composition of local maps, such as equation (53), should be suitably limited so that the composition is well defined.

### B. Higher-order Integrator

Using the first-order maps defined by (53) and its adjoint map

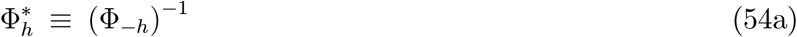

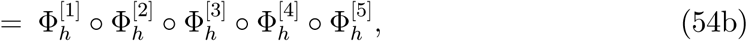

higher-order integrators can be obtained as a map by the symmetric composition with the adjoint [37]:

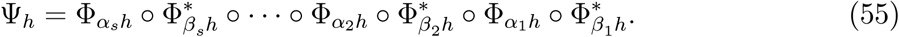

Here, coefficients {*α_i_*, *β_i_*} ⊂ ℝ satisfy the symmetric condition

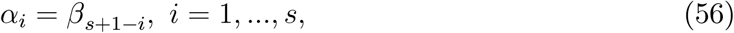

in order to satisfy the symmetric property
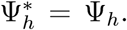
A number of specific values of the parameters, viz., stage s and coefficients {*α_i_*,*β_i_*}, are known [29, 37]. The simplest second-order integrator is a generalization of the Verlet method, defined by

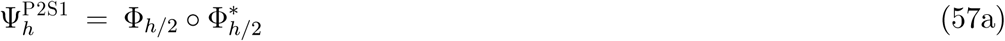

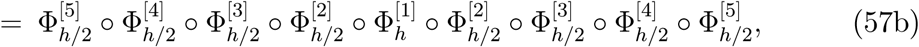

called as P2S1. Fourth-order integrators includes P4S5 [38] and P4S6 [39]. The integrator is thus explicit, symmetric (or time reversible: (*Ψ_h_*) ^−1^ = *Ψ*_–h_), and systematic, viz., higher-order integrators can be systematically constructed.

Note that the appearance ordering of
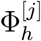(*j* = 1, …, 5) in equation (53) is arbitrary to ensure the first-order local accuracy. In contrast, the ordering has an influence on the computational time. The most time-consuming operand is the evaluation of the atomic force *F*(*x*) and its potential function *V*(*x*). The force evaluation is necessary for
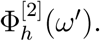 In addition, the evaluation of *F*(*x*) or *V*(*x*) may be required for
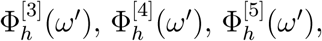
and invariant *L*(*ω*′), through *R*(*x*,*v*), *G*(*x*,*K*(*v*),*ζ*), *Y*(*ω*), and 𝓑(*ω*), respectively, depending on the definitions of these four quantities. For the typical form (57), the number of the force or potential evaluation is 1 in any cases if we use the ordering in equation (53), but otherwise it may be greater. Thus, one of the recommended ordering is given by (53). Since MD simulation spends almost all the calculation time for the evaluation of *F*(*x*) or *V*(*x*) and since it is often very heavy, less evaluation is highly required. In this sense, implicit integration methods are unfavorable and explicit methods like the current one would be advantageous. Ideas for increasing the efficiency of the integrator [40, 41] could be combined with the current method.

### C. Approximation

One possible problem in the current scheme is in seeking the specific form of 𝓚^*ω*^ (*h*)≡ 𝓚_*α*_(*h*) in equations (36). That is, there are situations such that (i) the integration for
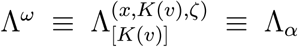
defined by (36b) cannot be explicitly represented by elementary functions; (ii) the integration for Λ^*ω*^ can be represented by elementary functions, but the inverse function 𝓚^*ω*^ = (Λ^*ω*^)^−1^ cannot be explicitly denoted for all h; (iii) 𝓚^*ω*^(*h*) can be explicitly denoted, but the domain of definition *J^ω^* = *J_α_* may be small in a practical use such that a fixed time step *h* may be beyond the range of *J^ω^* so that 𝓚^*ω*^ (*h*) is not well defined; (iv) even if 𝓚^*ω*^ (*h*) is safely obtained, it may be inconvenient to switch the maps, as represented in equation (51).

To avoid these situations, an approximation of Φ^[4]^ can be used. First consider the case [*K*(*v*)] ≠ 0. By noticing 𝓚^*ω*^ ∈ *C*^*r*+1^ (*r* ≥ 1) and *D*𝓚^*ω*^(*t*) = –2*G* (*x*,𝓚^*ω*^(*t*),*ζ*)(*t*)(see equation (37) or (43)) and assuming *G*(*x*,*k*,*ζ*) to be sufficiently smooth with respect to *k* (e.g., *r* ≥ 2), the Taylor expansion of *θ^ω^*(*h*) with respect to *h* around 0 yields

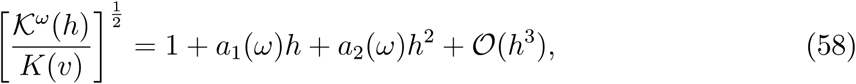

where

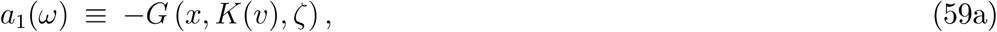

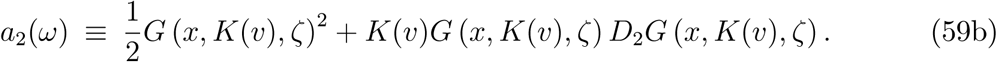

Thus

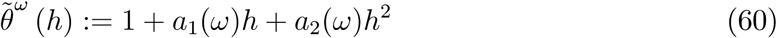

becomes an approximation of *θ^ω^*(*h*), when [*K*(*v*)] ≠ 0. For the case of [*K*(*v*)] = 0 (viz., *G* (*x*, *K*(*v*), *ζ*) *K*(*v*) = 0), a simple relation holds: *θ^ω^*(*h*) *v*= *v* = *θ͂^ω^* (*h*) *v* for any *h* and *ω*.

We thus have an approximation of Φ^[4]^, defined as

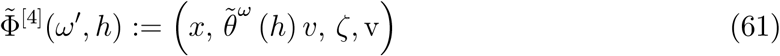

for all (*ω*′,*h*) ∈ Ω′ × ℝ, and

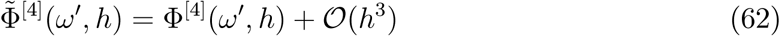

holds for all *ω*′ and *h*∈ *Ĵ^ω^*. Since the switching in equation (51) is vanished (or automatically involved) in equation (61), we have a solution to (iv) above. With this approximation, we also see that we do not encounter the situations (i)–(iii), so that the problems are solved.

Hence,
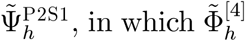
is used instead of
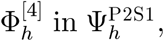
approximates 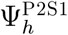 such that

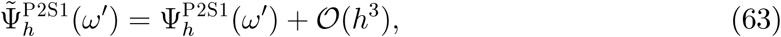

under a sufficient smoothness of the functions, *α*, *R*, *F*, and *G*. The time-reversible property also holds in the sense of this approximation. This indicates that utilizing Φ͂^[4]^ solves the problems but scarifies the exact time-reversible property. However, it should also be noted that these approximations become more accurate with a higher order expansion for equation (58), according to the smoothness conditions of the functions.

## V. APPLICATIONS

Several applications of the current method are demonstrated. According to the type of function *G* with respect to *k*, we here consider three cases and take into account examples of EOM to specifically construct Φ^[4]^.

Note that what we should do for *practically* using Φ^[4]^ for a *given* (initial value)*ω* = (*x*, *v*,*ζ*) is just to know 𝓚^*ω*^(*h*) and the specific representation of [*K*(*v*)] = 0 in equation (51). However, also for *theoretically* grasping the domain of definition of *h* for *all* (initial value) *ω* = (*x*, *v*, *ζ*) in advance, we need to know *J^ω^* in equation (50), for which it thus should not be
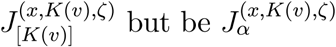
for every possible a; wherein we simply write as
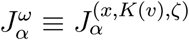
hereafter. Namely, regarding each type of *G*, what we shall do here is to know
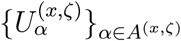
and the following three quantities for each *α* ∈ *A*^(*x*,*ζ*)^ satisfying
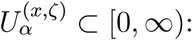

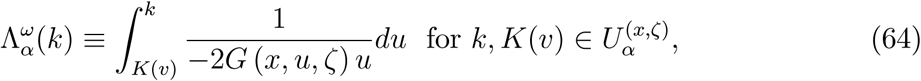

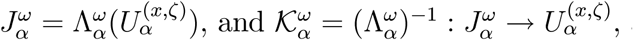 and finally find *θ^ω^* (*h*) for *h* ∈ *J^ω^*.

### A. Constant function

First, we consider a case in which *G*(*x*, *k*, *ζ*) is independent of *k* such that

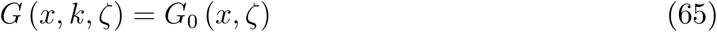

with a *C^r^* function *G*_0_: *D* × ℝ^*m*^ → ℝ. In this case, Φ^[4]^ is trivially obtained as

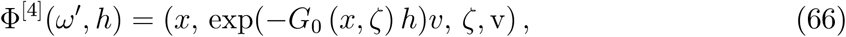

and we confirm below that this result is also obtained in the current formalism. As in section II, put *B* ≡ ℝ^*n*^ and *B*_K_ = ℝ. Then we see

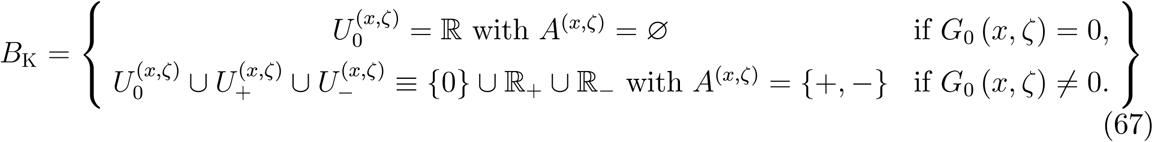

The condition 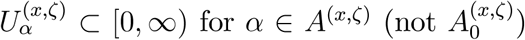 is only met by 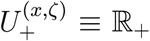 in the case of *G*_0_ (*x*_0_, *ζ*_0_) ≠ 0. In this case, we have

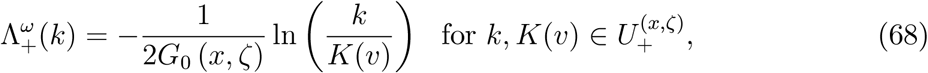

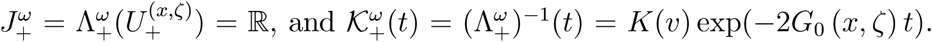 Since [*K*(*v*)] = 0 means *G*_0_(*x*, *ζ*) = 0 or *K*(*v*) = 0, we obtain

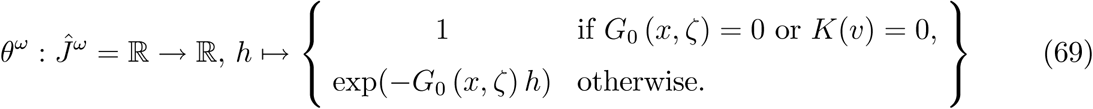

Finally, since *K*(*v*) = 0, *v* = 0, we recover equation (66) from equations (52) and (69).

The NH equation is one of the examples because of *G*_0_ (*x*, *ζ*) = *ζ*/*Q*, and the resulting map

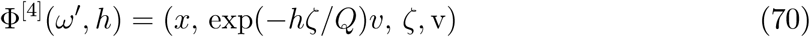

is essentially the one already obtained; see e.g., Ref. [27] in the context of the symplecticity for the original physical system and Ref. [25] in the context of the volume preservation [42, 43] for the total system. Note that the method can also be applied to the cNH lattice [44], which couples arbitrary numbers of the NH equations with distinct temperatures and has the simple friction form represented by equation (65).

### B. Laurent polynomial

We can treat *G* being a Laurent polynomial represented by

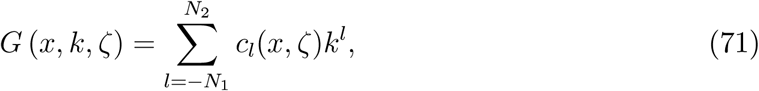

where *N*_1_ and *N*_2_ are non-negative integers and *c_ℓ_* are *C^r^* functions. In this case, since the integrand 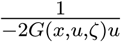 in equation (64) becomes a rational function with respect to *u*, the integral 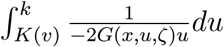 can be represented by elementary functions. Thus we can avoid situation (i) stated in section IV C. Even so, there may arise situations (ii)–(iv), requiring the approximation method. In the two specific examples we consider below, however, we do not encounter all situations (i)–(iv).

#### 1. The Berendsen EOM

The Berendsen EOM is one of the examples. As in section II, put 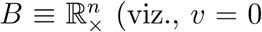 is forbidden), 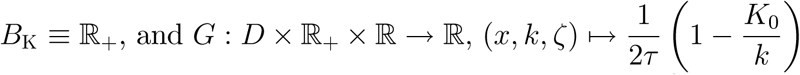 (viz., *N*_1_ = 1, *N*_2_ = 0; recall *K*_0_ is not the initial value but the EOM parameter). Then we can see that the decomposition of *B*_K_ is done as

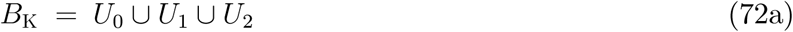

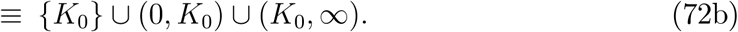

Note that since *G* is independent of (*x*, *ζ*), so are all quantities, 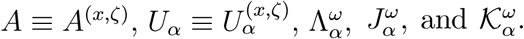 The condition *U_α_* ⊂ [0, ∞) for *α* ∈ *A* is obviously met by *α* = 1 and 2. In these cases, we have

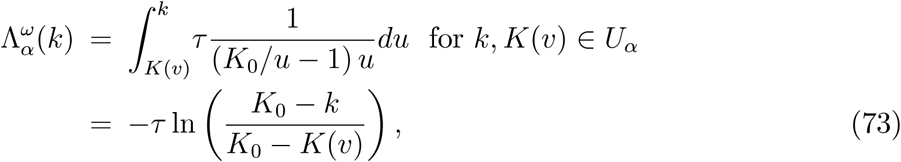

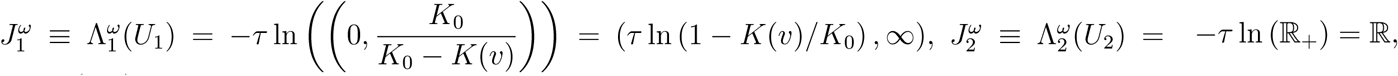 and

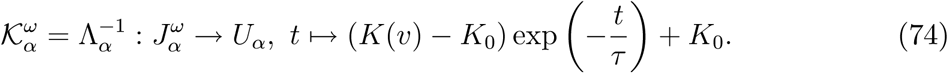

Since [*K*(*v*)] = 0 ⇔ *K*(*v*) = *K*_o_, we obtain

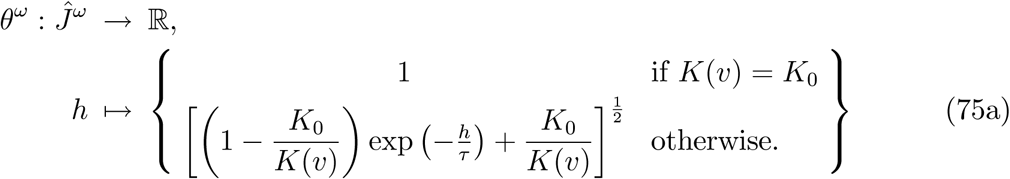

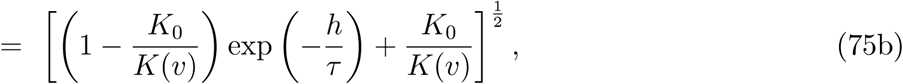

which eventually has a unified form without the switching. Hence, we do not encounter situations (i), (ii), and (iv). Regarding situation (iii), 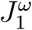 is not the whole ℝ, but 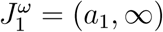 with *α*_1_ = **τ** ln (1 – *K*(*v*)/*K*_0_) < 0. However, 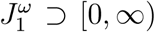 implies that any *h* ≥ 0 can be taken, so that there is no problem in a practical use (although a care may be needed to construct higher order integrators having negative coefficients *α_i_* and *β_i_*. Note that the resulting map

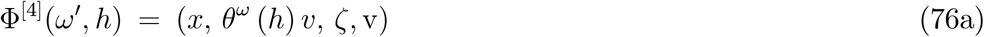

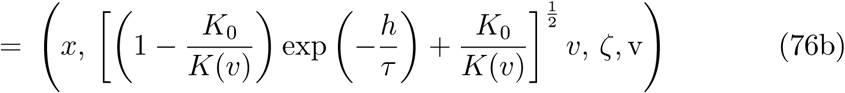

is obtained directly in Ref. [45]. We here thus arrived at the same result via a general theoretical procedure.

#### 2. The modified Berendsen EOM

As discussed in Ref. [46], another type of the Berendsen EOM,

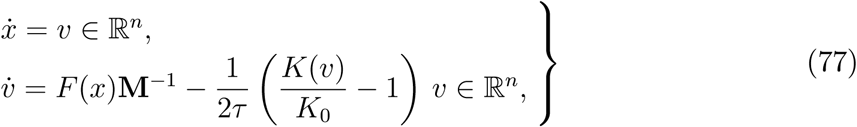

can be considered. Namely, instead of 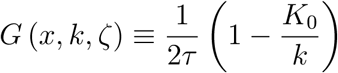 in the original Berendsen EOM, 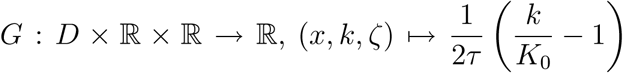 is used, which corresponds to the Laurent polynomial of the type of *N*_1_ = 0 and *N*_2_ = 1.

Put *B* ≡ ℝ^*n*^ and *B*_K_ = ℝ, where *v* = 0 is *not* forbidden. Similar to the original Berendsen EOM, independently of (*x*, *ζ*), we have the following decomposition:

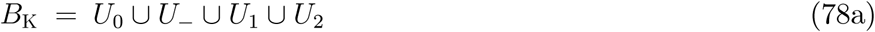

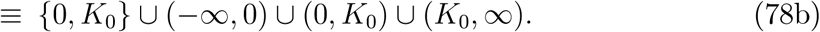

For the target *U_α_* ⊂ [0, ∞) (*α* = 1, 2), we have

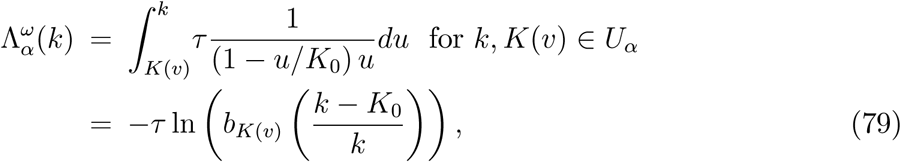

where *b*_*K*(*v*)_ ≡ *K*(*v*)/*K*(*v*) – *K*_0_),

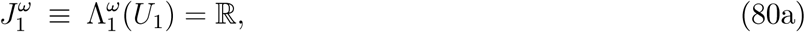

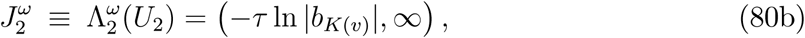

and

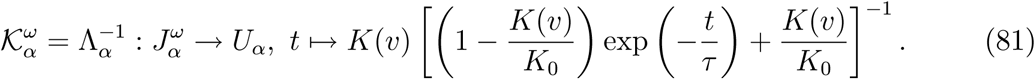

Since [*K*(*v*)] = 0 ⇔ *K*(*v*) ∈ *U*_0_ = {0, *K*_0_}, we get

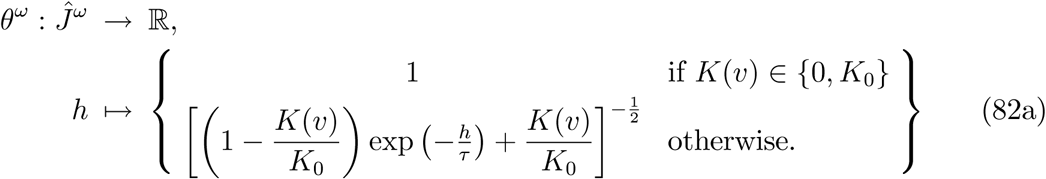

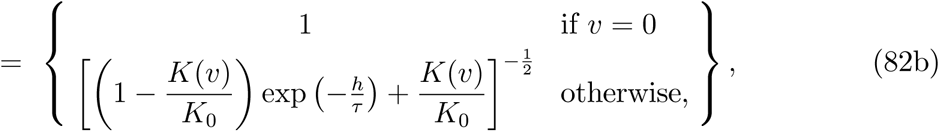

indicating a no-switching result,

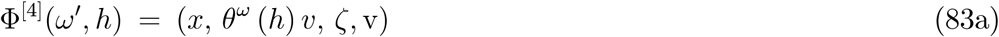

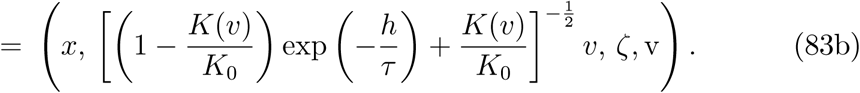

Hence, we do not encounter situations (i), (ii), and (iv) in section IVC. For situation (iii), although 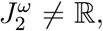 the fact that 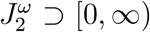 allows a use of any *h* ≥ 0. The resulted exact map (83b) is new, and it resembles equation (76b), as expected.

### C. Exponential function

We consider a case in which *G* (*x*, *k*, *ζ*) takes a form of

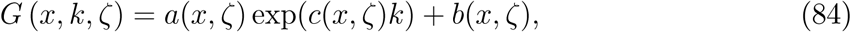

with *a*, *b*, and *c* being *C^r^* functions from *D* × ℝ^*m*^ to ℝ. Put *B* ≡ ℝ^*n*^ and *B*_K_ = ℝ. Then, according to the values of (*x*, *ζ*) ∈ *D* × ℝ^*m*^, there are several types of decomposition of *B*_K_. In any type of the decomposition, the condition 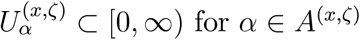 is met by one of the following types of intervals: (0, ∞), (0, *d*_(*x*,*ζ*)_), or (*d*(*x*,*ζ*), ∞), with a certain constant *d*(*x*,*ζ*) > 0. In these cases, the target quantity is

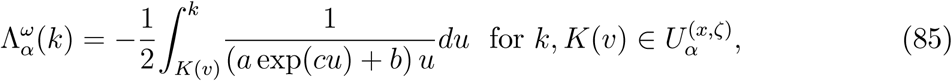

where *a* = *a*(*x*, *ζ*), *b* = *b*(*x*, *ζ*), and *c* ≡ *c*(*x*, *ζ*). In general, this integration cannot be represented with elementary functions; viz., situation (i) stated in section IVC occurs. Thus the approximation method in section IV C can be applied, and the approximated map Φ̃^[4]^ is given by equation (61) with equation (60), where *a*_1_(*ω*) and *a*_2_(*ω*) become

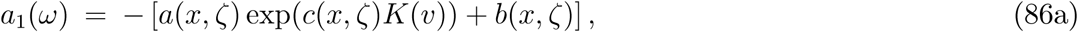

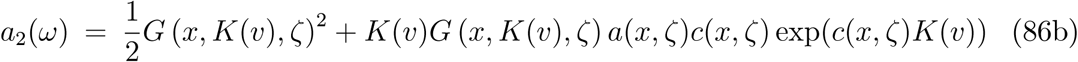

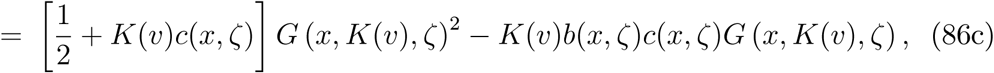

according to equation (59).

#### 1. The cNH EOM

As an example, we take the cNH defined in Examples 4 and 8. Since Φ^[4]^ for equation (7) is clear as also discussed in section V A, we consider that for equation (6). By choosing equation (22) for *ρ*_E_ with *n*_T_ = 1, the explicit form of 𝓣 is given by equation (27). In this case,

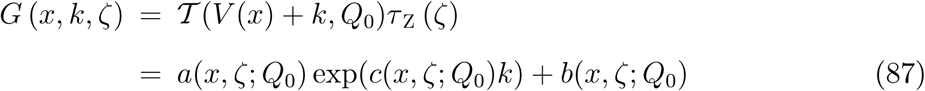

with

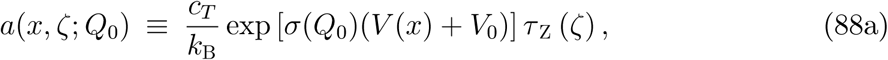

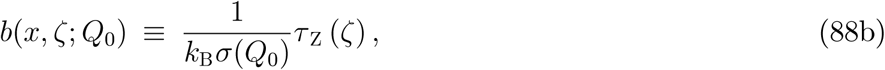

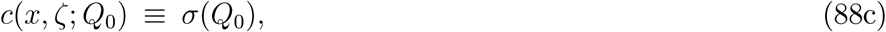

where *Q*_0_ dependence is also taken into account. Thus, the approximated map is given by

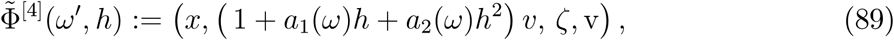

where

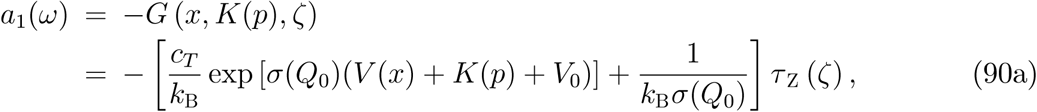

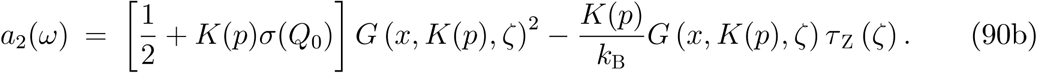

Note that only the case of *C_T_* = 0 was taken into account to integrate the cNH EOM in the previous work [30]. As discussed in section V A, *C_T_* = 0 is the easiest case since *G* becomes independent of *k* such that *G* (*x*, *k*, *ζ*,) = *τ_Z_* (*ζ*) /*k*_B_*σ*(*Q*_0_), and so Φ^[4]^ is easily obtained. Thus the above result is new and can be applied even for *C_T_* > 0, which results in a far complicated *G* form compared with that in the *C_T_* = 0 case.

## VI. NUMERICS

### A. Method

The three EOMs-NH, Berendsen, and cNH-have been numerically solved using the current integration method in order to confirm its ability. For simplicity, the most fundamental one, P2S1, defined in (57) has been used, where Φ^[*i*]^ (*i* = 1, 2, 3, 5) are defined in (29), with *α* and *R* being described in Examples in section II for each EOM. Φ^[4]^ are defined in equations (70) and (76) for the NH and the Berendsen, respectively, and its approximation Φ̃^[4]^ is defined in equation (89) for the cNH. The invariant functions defined by equations (17), (20), and (26), respectively, for the NH, Berendsen, and cNH, were monitored for confirming the accuracy of the integration.

For a physical system, one-dimensional harmonic oscillator (1HO) defined by the potential function,

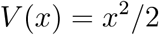

with *x* ∈ ℝ^1^ (viz., *n* = 1), was chosen, since it is the simplest model that describes the physical system behavior around an equilibrium state. The mass **M** =1 was used for all the three EOMs. All quantities are treated as dimensionless and *k*_B_ = 1.

The parameters and the initial values were set as follows. For the NH EOM, the target temperature *T*_ex_ was 1, the temperature-control constant was *Q* = 10^−1^. The initial values were *x*(0) = 1, *v*(0) = 1, *ζ*(0) = 0, and v(0) = 0. For the Berendsen EOM, the temperature-control time constant was *τ*=1 and the target kinetic energy was *K*_0_ = 1/2 (viz., the target temperature was 2*K*_0_/*nk*_B_ = 1). The initial values were *x*(0) = 1, *v*(0) = 1, and *v*(0) = 0. For the cNH EOM, the parameters used were **M**_T_ = 1 (*n*_T_ = 1), *c*_Z_ = *c*_Y_ = 1 [see equation (23)], and *c_T_* = 10^−6^ [see equation (27)]. Similar to the previous work [30], *ρ*_E_ was set by equation (22) with *V*_0_ = 0, *σ* was the sigmoid function [equation (70) of Ref. [30] with the parameters *β*_L_ = 0, *β*_R_ = 1, and *κ* = 10^−1^], and *f* was the Beta function [equation (72) of Ref. [30] with the parameters *p* = *q* = 5]. The initial values were *x*(0) = 1, *p*(0) = 1, *ζ*(0) = 0, *Q*(0) = 0, 𝓟(0) = 1, *η*(0) = 0, and v(0) = 0.

### B. Results

Results are shown in figure 1, together with figure S1 in the supplementary material, on the integration of the NH EOM with a unit time step *h* = 10^−3^. Figure 1 shows the behavior for a short time period (from *t* = 0 to 10^5^*h*) of the coordinate *x*, the system temperature *T*(*v*) ≡ 2*K*(*v*)/*nk*_B_ = *v*^2^, along with the trajectory of (*x*,*v*), and the invariant *L*(*ω*, v). The behavior of these quantities for a longer time period and the behavior for other quantities are shown in figure S1. The coordinate *x* and velocity *v* are rapidly oscillating (figures 1(a) and S1(a)) and they indicate almost periodic motion in a structured regular region in the phase space (figures 1(b) and S1(b)) [47, 48]. Against these oscillations, the invariant was well conserved (figure 1(c) and S1(c)), indicating that the numerical integration was successfully done. Temperature control was also good as observed by the averaged temperature 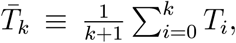 where *T_i_* = *T*(*v*(*ih*)), to be near the target temperature *T*_ex_ = 1, shown in figures 1(a) and S1(a), despite the fact that the BG distribution is not generated in the 1HO system due to the lack of the ergodicity [47].

**FIG. 1:**
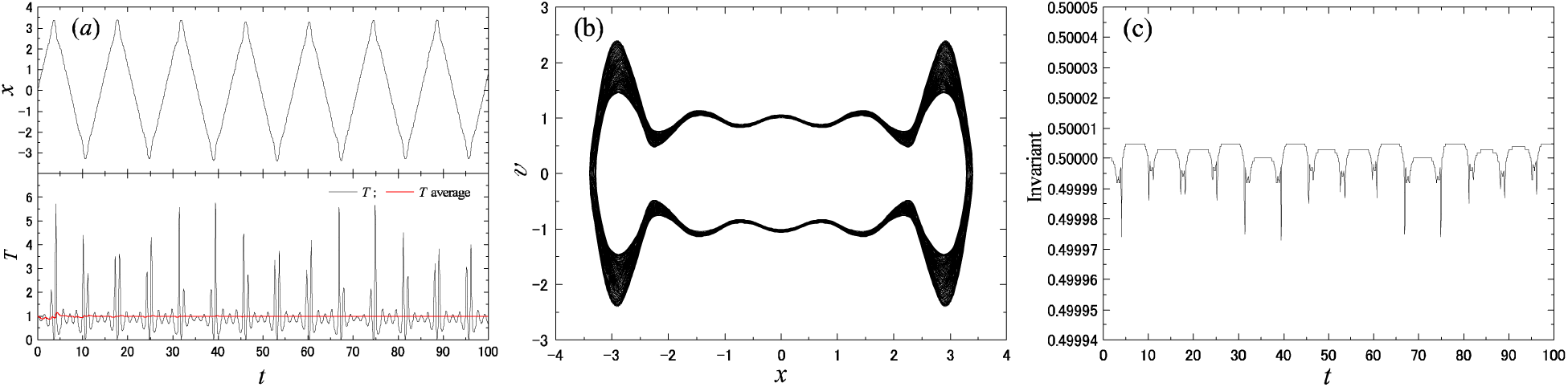
Results for the one-dimensional (1D) NH oscillator obtained by P2S1 for a short period of time *t*: (a) time development of coordinate *x*, system temperature *T* = *v*^2^, and the averaged temperature *T*; (b) trajectory of (*x*,*v*); and (c) time development of invariant *L*(*ω*, v).

In contrast, the Berendsen EOM indicates ill temperature control, as shown in figure 2(a), which is the result for *h* = 10^−4^. Monotonic decay of the temperature along with monotonic increase of the coordinate is observed. They are not due to the numerical error, since the integration was accurate as seen in the conservation of the invariant exhibited in figure 2(c). They are due to the special features of 1 dimensional system for the Berendsen EOM. That is, ODE (4) with *n* = 1 indicates that (i) there is no fixed point (since *v* = 0 is excluded); (ii) the (essential) phase space *D* × ℝ_×_ is disconnected and has two connected components Ω_±_ ≡ {(*x*,*v*) ∈ *D* × ℝ|*v* ≷ 0}, each of which is an invariant subspace (the extended phase space Ω⁢ also has two connected components), implying that any solution cannot go to the other half-plane by crossing the *v* = 0 line; (iii) in the upper half plane Ω_+_, the coordinate *x* always moves to the right direction because of *ẋ* = *v* > 0 and the prohibition of moving into Ω_–_(the converse holds in the lower half plane Ω_–_), implying that oscillating motion does not occur; and (iv) the (*x*, *v*) trajectory cannot cross with itself since the phase space is two dimensional, unlike the case of the NH EOM. These features can explain the qualitative behavior shown in figures 2(a) and (b). The vector field of the Berendsen 1HO system, (*x*, *v*) ↦ (*v*, –*x* + (*b*/*v*^2^ – *c*)*v*), shown in figure 3(a), can explain the quantitative behavior. These implies that the results obtained by the current integrator were correct.

**FIG. 2:**
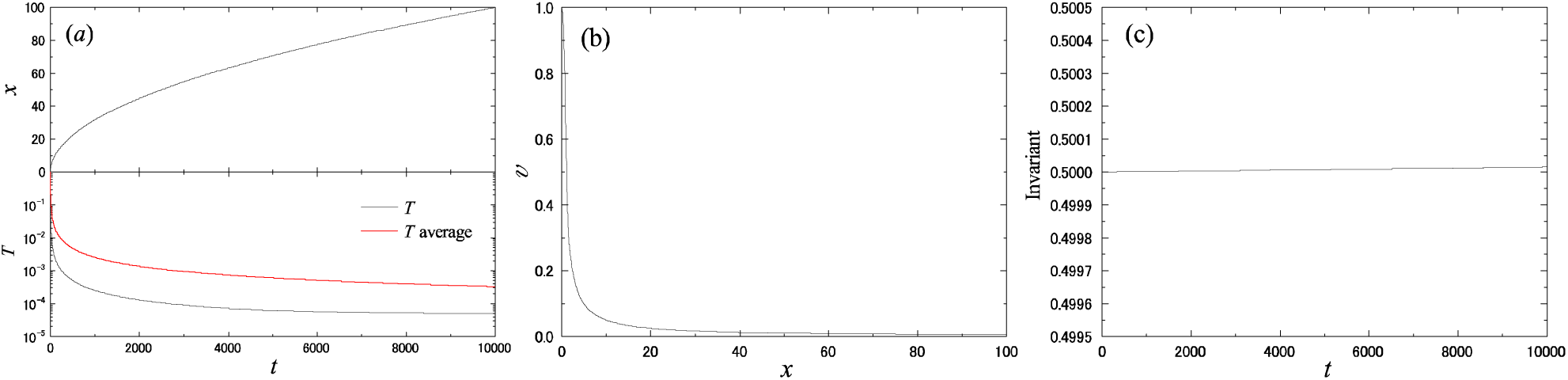
Results for the 1D Berendsen oscillator obtained by P2S1: (a) time development of coordinate *x*, system temperature *T* = *v*^2^, and the averaged temperature *T*; (b) trajectory of (*x*,*v*); and (c) time development of invariant *L*(*ω*, v).

**FIG. 3:**
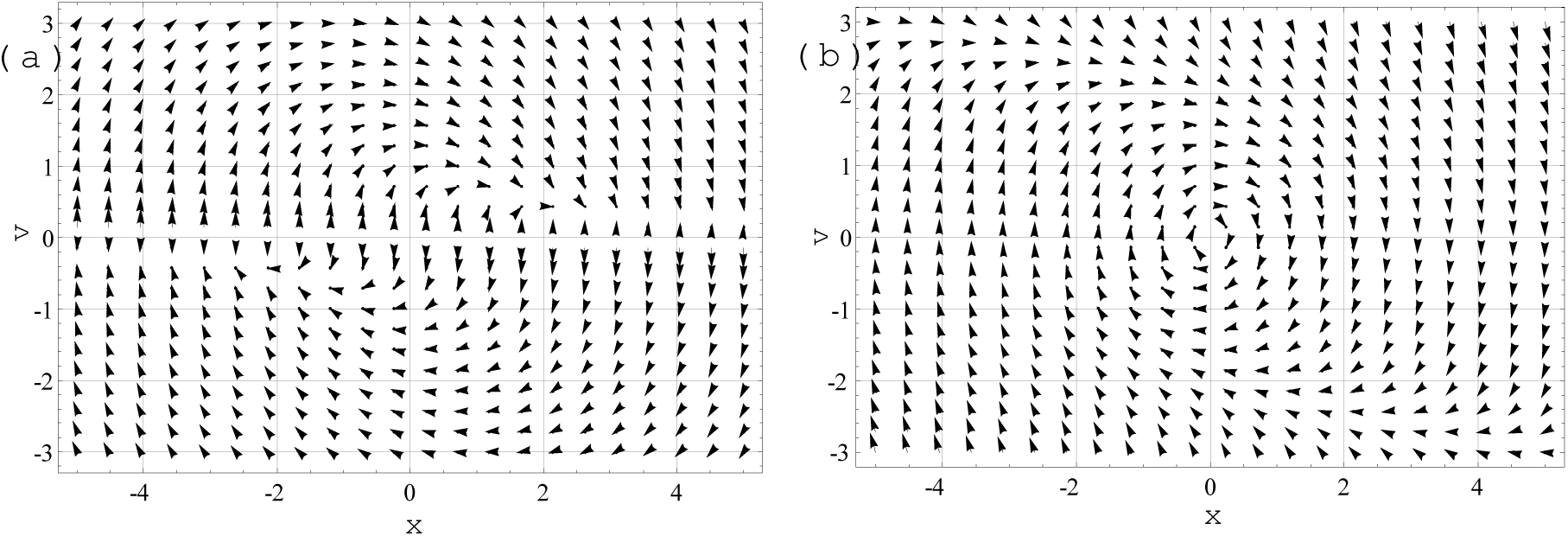
(a) Vector field of the Berendsen 1HO system, (*x*,*v*) ↦ (*v*, – *x* + (*b*/*v*^2^ – *c*)*v*), where *b* ≡ *K*_0_/**M***τ* = 1 and *c* = 1/2*τ* = 0.5. The prohibition of the crossing is due to the increase/decrease of the *v* component of the field –*x* + (*b*/*v*^2^ – *c*)*v* → ±∞ as *v* → 0±. The flow is point-symmetric about the origin, viz., *t* ↦ (–*x*(*t*), –*v*(*t*)) is a solution if *t* ↦ (*x*(*t*),*v*(*t*)) is a solution, which makes the 1-1 correspondence among the solutions in Ω_±_. (b) Vector field of the modified-Berendsen 1HO system, (*x*, *v*) ↦ (*v*, –*x* – (*dv*^2^ – *c*)*v*), where *d* ≡ **M**/4*τK*_0_ = 0.25 and *c* = 1/2*τ* = 0.5.

For comparison, results obtained by the conventional numerical integrator of the Berendsen EOM [4] are shown in figure 4, where the simulation conditions were the same as those stated above. The prohibition of the crossing is broken, as seen in figure 4(b), and it occurs sudden and periodically as seen in figure 4(a). These results were derived via the specific feature of the “stiff” vector field and the numerical error, which can be confirmed in figure 4(c). Integration of this system with a high accuracy and low cost is tough in general in that it has two sets closely located each other, one is a “stable” set, which is supposed as seen in figures 2(b) and 3(a), and the other one is the boundary ℝ × {0}, which has a strong “repulsion”. At any rate, the error of the conventional scheme was significant judging from the invariant non-conservation.

**FIG. 4:**
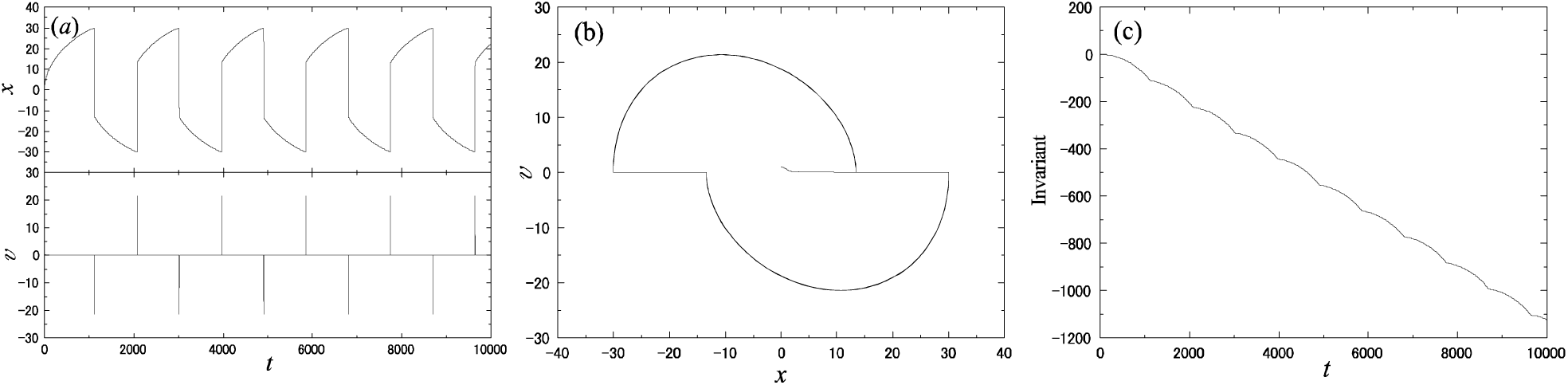
Results for the 1D Berendsen oscillator obtained by the conventional method: (a) time development of coordinate *x* and velocity *v*; (b) trajectory of (*x*,*v*); and (c) time development of invariant *L*(*ω*, v).

1HO was investigated here since the same fundamental physical model should be used for the three EOMs in comparison. Note also that the purpose of this study is not to judge the performance of each EOM, such as the temperature controllability, but is to examine the properties of the integrators for given ODEs. However, 1HO nevertheless seems peculiar for the Berendsen EOM, as stated. In fact, for larger systems (the phase space is connected for *n* > 1), the Berendsen EOM works well to conduct the temperature control of the physical system (see e.g., Ref. [49]). On the other hand, the integrator accuracy is also important for the temperature controllability even in larger systems [45]. Note also that 1HO does not seem so peculiar for the modified Berendsen EOM introduced in section VB 2. Since its vector field, (*x*, *v*) ↦ (*v*, –*x* – (*dv*^2^ – *c*)*v*), has a unique fixed point (0, 0) (see figure 3(b)) and since the phase space is connected (*D* × ℝ = ℝ^2^), the behavior should be the one perturbed from that of the Newtonian EOM. The expected results of the modified Berendsen EOM are exhibited in figure S2.

Figure 5 shows the results for the cNH EOM obtained by the current integrator with *h* = 10^−3^. Fluctuations of larger amplitude *p* = *v* are observed, compared with that in the NH EOM, as expected from the feature of this EOM, viz., the heat bath temperature (which corresponds to *T*_ex_ in the NH EOM) fluctuates according to the predetermined distribution function. Following the fluctuations of *p*, the coordinate *x* also shows larger fluctuations. The invariant, nevertheless, almost keeps constant, showing a robustness of the current integration scheme. The results for the other quantities are shown in figure S3. Figure S4 indicates that the conservation of the invariant in the current integrator was better for a smaller *h*, which is expected, and the behavior of *x* and *p* remains similar, which is not so expected considering the chaotic feature of this system.

**FIG. 5:**
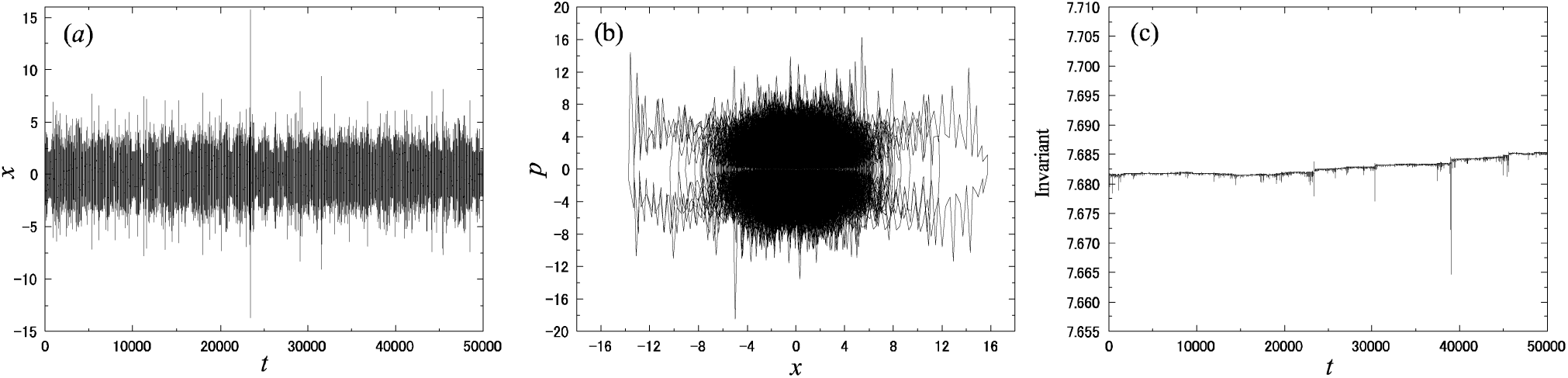
Results for the 1D cNH oscillator obtained by P2S1: (a) time development of coordinate *x*; (b) trajectory of (*x*; *p*); and (c) time development of invariant *L*(*ω*; *v*).

Figure 6 shows the results for the cNH EOM obtained by the conventional 4th order Gear integrator. Larger fluctuations of *p* and *x* were generated as in the current P2S1 integrator, but the conservation of the invariant was not good and rapidly changed. On the other hand, the results of the Gear integrator with a smaller *h* shown in figure S5 halfway keeps the accuracy that is higher than that of the current method. One of the reasons is that the Gear method has the 4th-order local accuracy while the current method 2nd order. Note that, however, after then, a sudden jump, which should be due to large fluctuations of variables, also occurred in the Gear integrator even with this smaller h. These results indicate that the current integrator outperforms the general purpose 4th-order Gear integrator. Note also that twice evaluations of the potential function *V*(*x*) are needed in the Gear integrator to monitor the invariant function defined by equation (26), while the evaluation is once in the current P2S1 integrator.

**FIG. 6:**
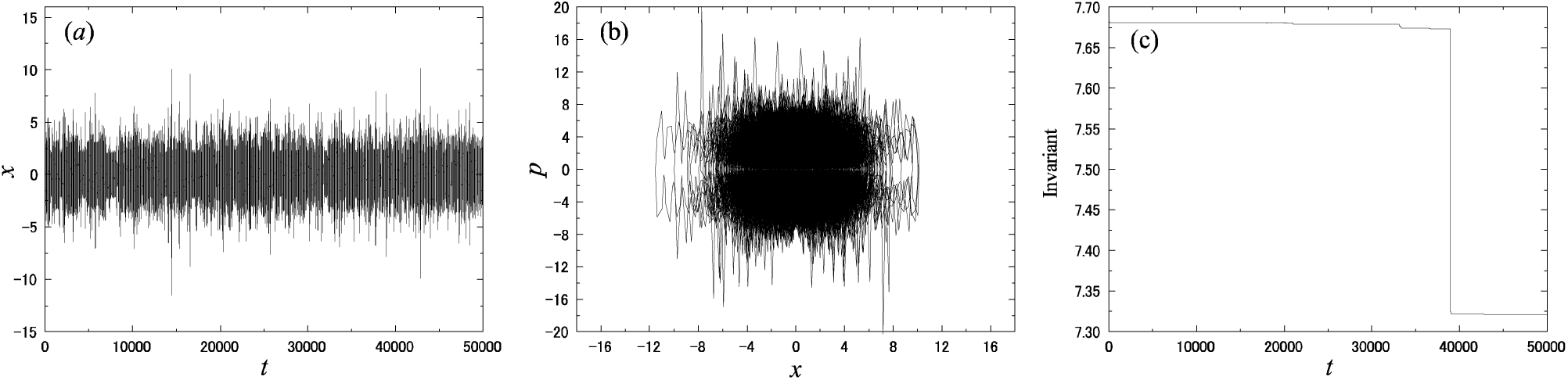
Results for the 1D cNH oscillator obtained by the Gear method: (a) time development of coordinate *x*; (b) trajectory of (*x*; *p*); and (*c*) time development of invariant *L*(*ω*; *v*).

## VII. CONCLUSION

A general scheme to construct symmetric, explicit numerical integrators for molecular dynamics equations of motion with a generalized friction is provided. Its theoretical discussion in depth is the main purpose of this study. The generalized friction is assumed to be the form of – *G*(*x*,*K*(*v*),*ζ*) *v*, which is proportional to the atomic velocity vector *v* ∈ ℝ^*n*^ scaled with the friction “constant” *G* (*x*,*K*(*v*),*ζ*) depending on the kinetic energy *K*(*v*) = (*v*|*v***M**)/2. The provided scheme is based on an exact solvable decomposition of the vector field of the target ODE, where the key point is to solve equation (30), which is essentially *v̇* = –*G* (*x*, *K*(*v*),*ζ*) *v*. Although this equation is multidimensional, it intrinsically has a one-dimensional structure to be exactly solved. Under a general condition, this has been performed, yielding a compact form of the phase-space map. The domain of definition of map obtained for each vector field is also exactly described. In addition, to avoid possible technical problems and reduce inconveniences in a practical use, an approximation method is also presented. Among the integrators provided here, the second-order P2S1 integrator should be robust for large variation of variables and suitable for MD since the evaluation of atomic force and potential is once at each time step.

As prototypical applications, the NH EOM, the Berendsen EOM, and the cNH EOM have been considered. Although the complicatedness of these ODE expressions differ, the integration scheme framework completely works for the all. For numerical check, the simplest but effective P2S1 integrator was applied to the 1HO model for the three EOMs. The solutions obtained by the current initial conditions and the EOM parameter values were almost periodic for the NH EOM, monotonic for the Berendsen EOM, and chaotic for the cNH EOM. Judged from the conservations of the invariant functions, the integrator was sufficiently accurate.

Regarding a deeper investigation of each EOM should be a separate topic. For the NH EOM, the results obtained here are not new, but the theoretical formalism provided here should be used for EOMs that have similar mathematical structures but have different physical background. Studies of parameter dependence, robustness, and time reversibility of integrators for the Berendsen EOM are studied in Ref. [45]. The current Berendsen integrator also used there has been theoretically derived in the current paper for the first time and considered it in a uniform mathematical ground of a generalized friction. Although the Berendsen 1HO presents a peculiar model to be physically less important, it may provide a mathematically interesting model for basic numerical integration study. The modified Berendsen EOM has not been studied much in the literature, but it is tractable and should have good properties to be investigated for more details. Ergodic property of the cNH studied in [15, 16, 30] is theoretically critical for ensuring the realization of the density and practically important as a phase-space sampling method. On the basis of the current basic study, applications to larger systems are under the study.

Last, additional remarks on general utilities of the method are made. The scheme provided here can be used for EOMs other than the three examples, and it may be used for other purpose than the temperature controlling, distributing, and fluctuating. The techniques may de translated into a scheme for treating the pressure of the physical system and beyond the molecular dynamics context. Higher-order integrator, such as fourth-order methods P4S5 and P4S6, can also be used, which is useful to attain higher accuracy for a small time step. The proposed invariant function defined on the extended phase space is simple and useful for any integrator to detect its numerical errors. Although this enjoys the fact that the EOM is represented by an ODE, the current technique could also be used for stochastic differential equations, where the error estimate would need much effort in general, as detailed by the analysis of the systematic bias of the splitting method especially for the Langevin dynamics for the BG distribution sampling [50].

## ACKNOWLEDGMENTS

This work was supported by a Grant-in-Aid for Scientific Research (C) (25390156 and 17K05143) from JSPS and the “Development of core technologies for innovative drug development based upon IT” from Japan Agency for Medical Research and development, AMED.

